# A 3-fold kernel approach for characterizing Late Onset Alzheimer’s Disease

**DOI:** 10.1101/397760

**Authors:** Margherita Squillario, Federico Tomasi, Veronica Tozzo, Annalisa Barla, Daniela Uberti, for the Alzheimer’s Disease Neuroimaging Initiative

## Abstract

The purpose of this study is to identify a global and robust signature characterizing Alzheimer’s Disease (AD). Two public GWAS datasets were analyzed considering a 3-fold kernel approach, based on SNPs, Genes and Pathways analysis, and two binary classifications tasks were addressed: cases@controls and APOE4 task. In the SNP signature of the ADNI-1 and ADNI-2 datasets, chromosome 19 and 20 reached high classification accuracy. In addition, the functional characterization of ADNI-1 and ADNI-2 SNP signatures found enriched the same pathway (i.e., Neuroactive ligand-receptor interaction), with GRM7 gene in common with both. TOMM40 was confirmed linked to AD pathology by SNP, gene and pathway-based analyses in ADNI-1. Using this 3-fold kernel approach, a peculiar signature of SNPs, genes and pathways has been highlighted in both datasets. Based on these significant results, we retain such approach a valuable tool to elucidate the heritable susceptibility to AD but also to other similar complex diseases.

## 1. Introduction

Alzheimer’s disease (AD) is the predominant form of dementia (50-75%) in the elderly population. Two forms of AD are known: an early-onset (EOAD) that affects the 2-10% of the patients and is inherited in an autosomal dominant way, with the three genes APP, PS1 and PS2 mainly involved; and a late-onset form (LOAD) that affects the vast majority of the patients in the elderly over 65s, whose causes remain still unknown (Van Cauwenberghe, Van Broeckhoven and Sleegers, 2016). Although LOAD has been defined as a multifactorial disease and its inheritance pattern has not been clarify yet, it is coming out the idea that it could be likely caused by multiple low penetrance genetic variants (Naj, Schellenberg and Alzheimer’s Disease Genetics Consortium (ADGC), 2017), with a genetic predisposition for the patients and their relatives estimated of nearly 60-80% (Naj, Schellenberg and Alzheimer’s Disease Genetics Consortium (ADGC), 2017).

The first well known gene associated to LOAD was APOE (Pericak-Vance *et al.*, 1997). It encodes three known isoforms proteins (ApoE2, ApoE3 and), with APOE4 known to increase risk in familial and sporadic EOAD. This risk is estimated to be 3-fold and 15-fold for heterozygous and homozygous carriers respectively, with a dose-dependent effect on onset age (Naj, Schellenberg and Alzheimer’s Disease Genetics Consortium (ADGC), 2017).

Large-scale collaborative GWAS and the International Genomics of Alzheimer’s Project have significantly advanced the knowledge regarding the genetics of LOAD (Van Cauwenberghe, Van Broeckhoven and Sleegers, 2016). Anyways, none of the new identified loci reached the magnitude of APOEε4, as predisposing risk factors for AD, with the majority of the hereditable component of AD remaining unexplained (Gandhi and Wood, 2010). Several different but not mutually exclusive explanations of such failure could coexist: AD could be caused by the concerted action of independent genetic factors, each having a small effect size that require to adopt multivariate methods and increased sample size (Moore, Asselbergs and Williams, 2010); or it could be caused by the concerted actions of multiple genes (again characterized by low effect size) that act inter-dependently in still undefined pathways, that would need a pathway-based approach, as done for other complex diseases (Frazer *et al.*, 2009). Alternatively, AD could be caused by vary rare but highly penetrant mutations that might be identified through DNA sequencing (Ng *et al.*, 2008).

In order to explore the first two possible scenarios, in this study we proposed a GWAS analysis based on multivariate methods and on a **telescope** approach, in order to guarantee the identification of correlated variables, and reveal the possible connections existing among the identified relevant variables at the different, but biologically connected, SNPs, genes and pathways levels.

Figure 1 depicts the workflow that we defined “3-fold kernel approach”: the term *3-fold* underlines the analyses at the SNP, gene and pathway level, and the word “*kernel”* is a synonymous of machine learning methods. The final purpose is to identify lists or signatures of possible causal SNPs, genes and pathways that considered together might provide a convincing picture of heritable factors in the LOAD pathogenesis.

**Fig. 1.**
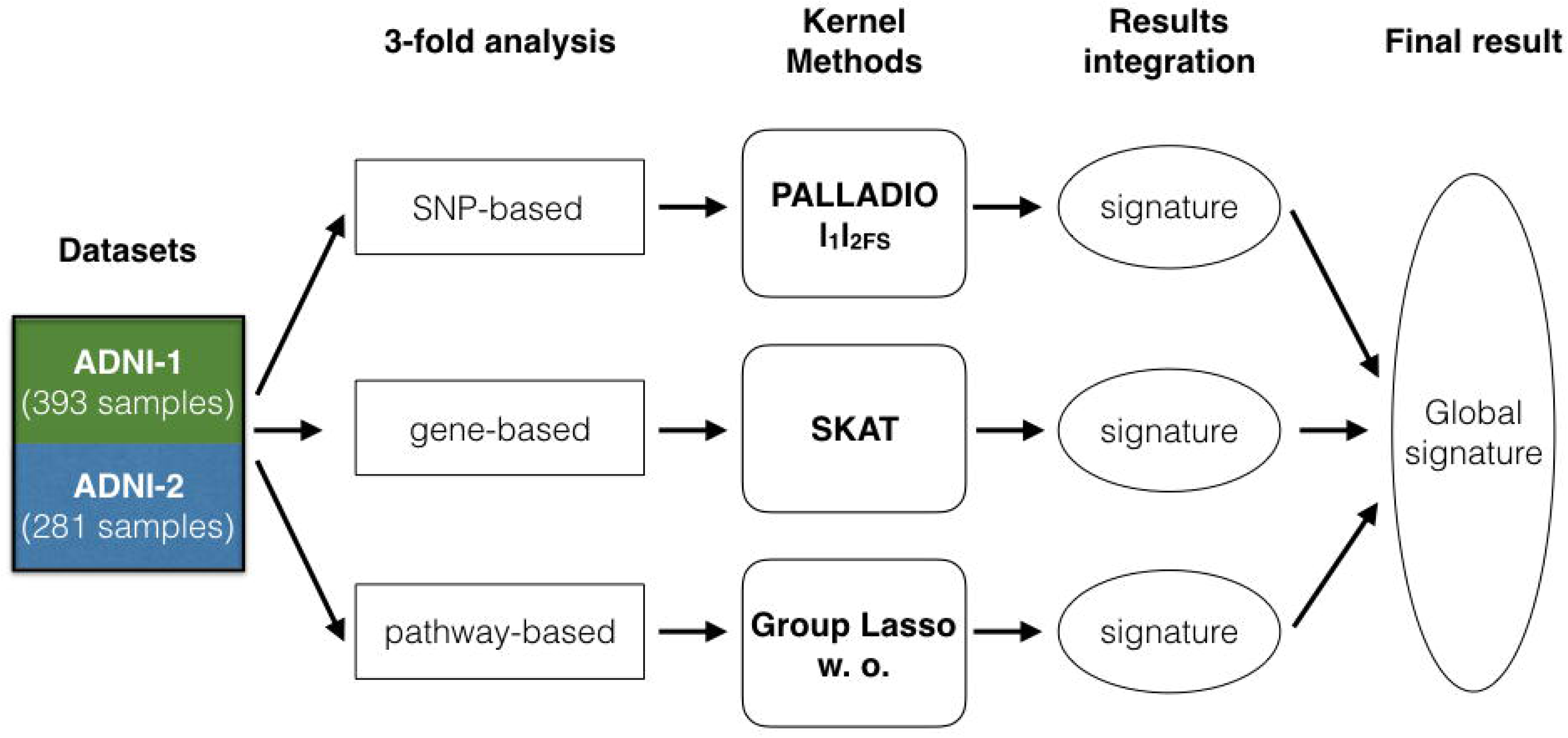
Workflow of the study. ADNI-1 and ADNI-2 datasets were analyzed at the SNP, gene and pathway level using three kernel approach with machine learning methods. The global signature represents the summary of the single integrated signatures identified in the 3-fold analyses.

## 2. Results

### 2.1 SNP-based analysis

SNP-based analysis performed on ADNI-1 dataset identified a signature of 14 SNPs relevant for cases@controls task (Fig. 2 and Table S3). These SNPs, mapped on 14 genes or intergenic regions, are located on chromosomes 6 and 2/0. In particular, chromosome 6 showed higher performance values, considering both balanced accuracy and MCC (0.61± 0.06 and 0.21±0.13) (Fig. 2A). In addition, the higher distance between the regular (light blue) and the permutation (red) distributions of the calculated balanced accuracies reinforced the robustness of the obtained results. While for chromosome 20, the partial overlapping between the regular and permutation batch gave less reliability, still maintaining a good performance in term of balanced accuracy and MCC (0.55±0.06 and 0.11±0.12) (Fig.2B).

**Fig. 2.**
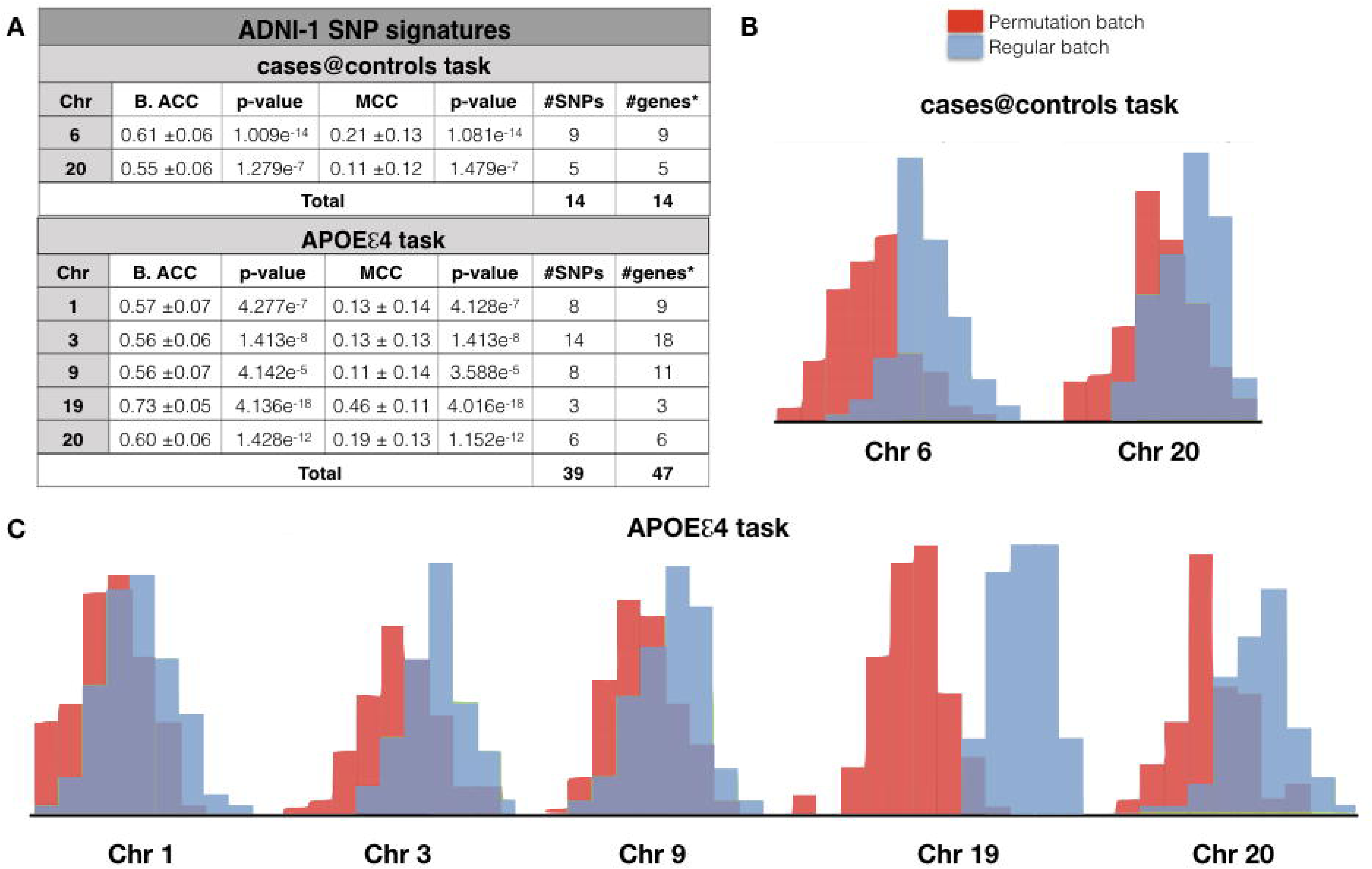
SNP-based results of ADNI-1. **A)** The classification performance of SNP based analyses performed in ADNI-1 considering two classification tasks: AD vs. healthy controls (cases@controls) or 1/2 APOE ε4 vs. 0 APOE ε4 carriers (APOEε4 task). B. ACC, Balanced Accuracy; MCC, Matthews Correlation Coefficient; #genes*, number of genes or intergenic regions**. B**) Balanced accuracy distribution plots of the regular (light blue) and the permutation batches (red) related to chromosomes 6 and 20 in the cases@controls task. **C**) Balanced Accuracy distribution plots of the regular (light blue) and the permutation (red) batches related to chromosomes 1, 3, 9, 19 and 20 in the APOEε4 task.

It is well recognized that *APOE* polymorphic alleles are the main genetic determinants of AD risk, being the individuals carrying one or two ε4 alleles at higher risk to develop AD (Pericak-Vance *et al.*, 1997). Thus, a further SNP based analysis was performed basing on the binary classification 1 or 2 APOEε4 vs 0 APOEε4 presence (APOEε4 task). 39 SNPs, which map to 47 genes or intergenic regions, have been identified in the APOEε4 task (Fig. 2A and Table S3). Chromosomes 19 and 20 were associated with the highest balanced accuracy and MCC results (Fig. 2A) and the distribution plots underlines this result (Fig. 2C).

Interestingly, the two classification tasks had in common the 20orf196 gene on chromosome 20. This gene is the closest to different SNPs found discriminant in the two tasks: in cases@controls task **rs6053572** is located in the intergenic region between GPCPD1 and 20orf196 while in the APOEε4 task **rs236137** and **rs6041271** are located in the intergenic region between 20orf196 and CHGB (Table S3).

SNP-based analysis on ADNI-2 dataset identified for cases@controls task a signature of 138 SNPs, which map to 183 genes or intergenic regions harbored on 19 different chromosomes, with a balanced accuracy and MCC values ranging from 0.63 to 0.81 and 0.26 to 0.63 respectively (Fig. 3A and Table S4). In particular chromosomes 9, 10, 14, 20 and 21 are the most reliable since they showed a higher distance between the two distribution measures (Fig. 3B).

**Fig. 3.**
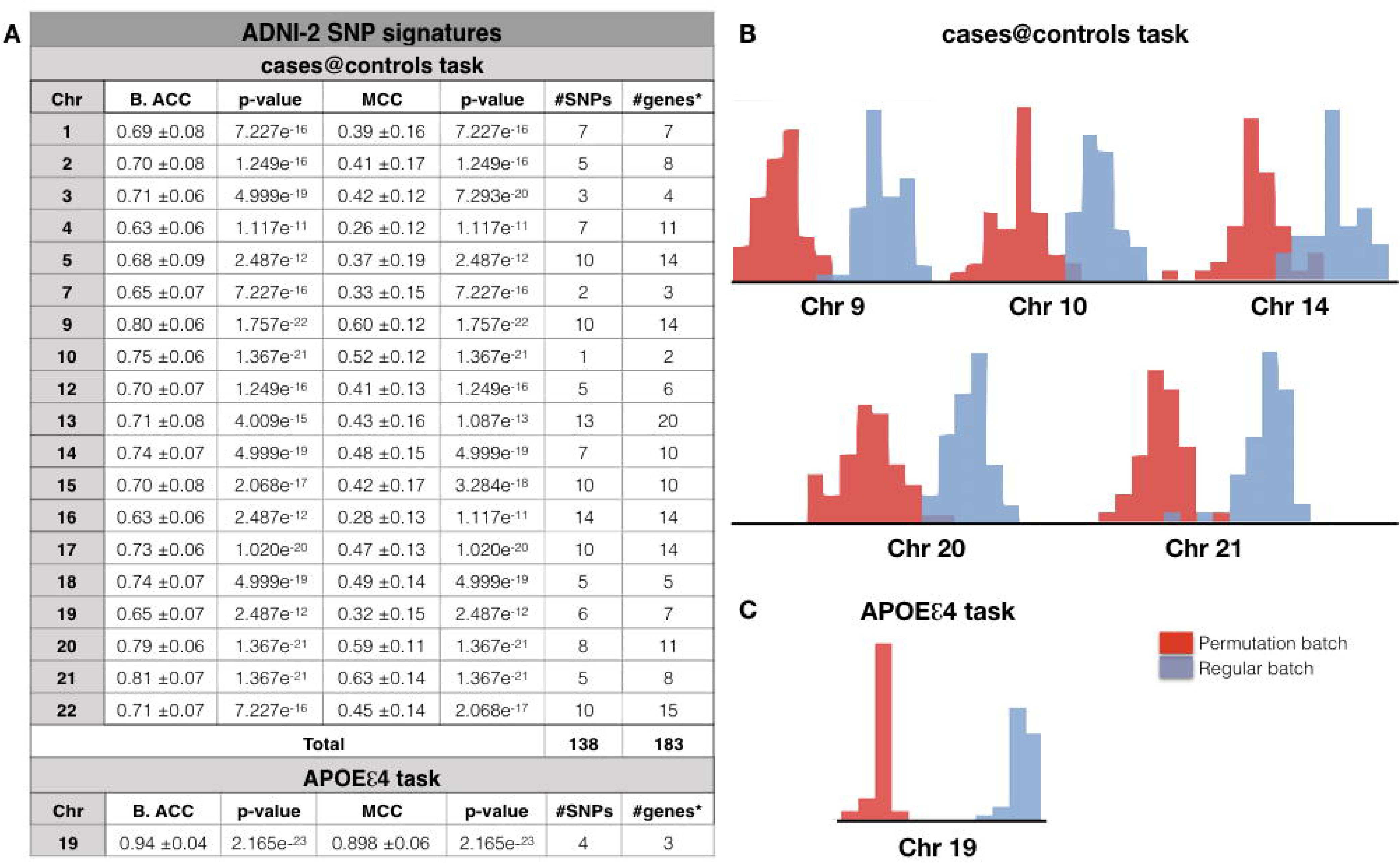
SNP-based results of ADNI-2. **A)** The classification performance of SNP based analyses performed in ADNI-1 considering two classification tasks: AD vs. healthy controls (cases@controls) or 1/2 APOE ε4 vs. 0 APOE ε4 carriers (APOEε4 task). B. ACC, Balanced Accuracy; MCC, Matthews Correlation Coefficient; #genes*, number of genes or intergenic regions**. B**) Balanced accuracy distribution plots of regular (light blue) and permutation (red) batches related to chromosomes 9, 10, 14, 20, 21 in the cases@controls task. **C**) Balanced accuracy distribution plots of regular (light blue) and permutation (red) batches related to the chromosome 19 in the APOEε4 task.

When we considered the APOEε4 task, only chromosome 19 was found statistically significant with very high values of both balanced accuracy (0.94) and MCC (0.90) (Fig. 3A and Table S4) and with very high distance between the two distributions (Fig. 3B). The derived SNP signature harbored only four SNPs located in three genes: **rs367209** in LOC101928063, **rs383133** in ZNF221, **rs415499** in ZNF155 and **rs365745** that causes a missense mutation in ZNF221 gene. None of these genes are known to be associated to AD but the AlzGene database (Bertram *et al.*, 2007) confirmed that these SNPs are located in a linkage region (*i.e.*, 19q13.31) known to be associated to AD.

### 2.2 Functional characterization of SNPs signature

In order to biologically characterize the gene lists derived from the SNPs signatures, we further performed a functional characterization in KEGG and in DISEASE database (see Supplementary Materials) for both datasets (Table 1). For ADNI-1 dataset, only APOEε4 task SNP signature was used, since cases@controls task did not allow this kind of analysis for the shortness of SNPs list identified. Interestingly, only the “**Neuroactive ligand-receptor interaction**” pathway, that includes the genes P2RY13, **GRIN3A**, LEPR, **GRM7**, P2RY14, reached a significant adjusted P value (Adj-P value = 5.99e-06) (Table 1). In addition, the enrichment in the DISEASE database highlighted the association of CHGB, **TOMM40**, **APOC1** and NGF genes with AD (Table 1). On the other hand, others diseases were found associated with the list of genes related to the ADNI-1 SNP signature: in particular, CHGB, TOMM40, APOC1 and NGF were in common with Tauopathies; GRIN3A, and KALRN genes were found associated with Schizophrenia; GRM7 was found associated to both Bipolar Disorder and Schizophrenia (Table 1). Other examples of diseases categories that reached significance (P value <.05) were mental disorders, dementia, and eating disorders, suggesting that the same genes might participate in the heritable susceptibility of different pathological conditions.

In ADNI-2 the functional characterization in KEGG and in DISEASE databases was allowed only for the cases@controls SNP signature, that reported a long SNPs’ list. From this analysis the most important pathways related to AD were: “**Chemokine signaling**”, “**Calcium Signaling**”, “**Axon Guidance**” and “**Neuroactive ligand-receptor interaction**” (Table 1). The latter pathway involved **GRIN2A**, **GRM7**, GABRG3 and CYSLTR2 genes. The result of the enrichment in DISEASE highlighted several relevant diseases or disease categories such as Bipolar Disorder, mood disorders, anxiety disorders, Schizophrenia (Table 1).

Bipolar Disorder, mood and eating disorders, Schizophrenia, nervous system diseases and mental disorders were in common with those found in the ADNI-1 functional characterization (APOEε4 task). Interestingly, Bipolar Disorder was the disease most enriched in ADNI-2, harboring ten genes (GRM7, GRIN2A, RBFOX1, GABRGB, PCDH17, ADCY9, PLCB4, SORCS2, AGAP1, MPPE1) of the cases@controls SNP signature.

### 2.3 Gene-based analysis

In order to identify a gene signature for cases@controls and APOEε4 tasks, ADNI-1 and ADNI-2 datasets were analyzed by using three different tests included in the SKAT software (see Methods). In ADNI-1 dataset, **TOMM40**, with its three SNPs, rs2075650, rs157580 and rs8106922, was found significantly associated to AD, applying all the tests; while TEF gene, harboring the three SNPs, rs738499, rs2073167 and rs17365991, was found significant in distinguishing cases@controls only with SKAT test (Table 2). When the ADNI-1 dataset was analyzed considering the APOEε4 task, we obtained very similar results using the three tests (Table 2). In particular, the genes or intergenic regions found significantly associated with the AD risk were: TOMM40, the intergenic region between LOC100129500 and **APOC1**, and the intergenic region between TOMM40 and APOE. Specifically three SNPs were located in the introns of TOMM40, the same found in the cases@controls task, while other two and one SNPs were located in the two just mentioned intergenic regions.

In the analysis of ADNI-2 dataset, considering both classification tasks, no genes or intergenic regions reaching the significance level were found (data not shown).

### 2.4 Pathway-based analysis

In REACTOME database (Fabregat *et al.*, 2018) we selected 9 pathway groups (Table S2), whose relevance in neurodegenerative processes were well recognized (Rosenthal and Kamboh, 2014) and we analyzed the SNPs mapped to the gene lists involved in these pathways. With ADNI-1_cases@controls task no groups reaching statistical significance were found. At variance, different pathway groups, associated with AD risk (APOEε4 task), achieved a good test score. These pathways and their relative groups were reported in Table 3.

Also in ADNI-2 dataset, several significant pathways associated with both classification tasks were identified (Table 3). Interestingly, “cellular senescence” and “detoxification of reactive oxygen species” pathways included in group 1c were in common between the two classification tasks. In addition, the “detoxification of reactive oxygen species” pathway of 1c group was also in common with the ADNI-1 pathway signature, while groups 5a and 9a of the ADNI-2_APOEε4 task were in common with ADNI-1_ APOEε4 task.

## 3. Discussion

Despite the promise of GWAS to reveal the genetic contribution to AD susceptibility, the majority of its heritable component remains unexplained. The major factor contributing to hamper the identification of genetic burden, lies in the cumulative contribution of multiple genes of small effect size that act inter-dependently in unidentified pathways. In addition, the need to adopt highly stringent statistical correction to avoid false-negative and false-positive results requires to increase the number of cases/controls in the initial study and then to reproduce the association across independent replicative studies. This study showed a new approach to analyze GWAS datasets in order to contribute in uncovering a robust and global AD signature. Here we discussed the data obtained by SNP, gene and pathway based analysis of two ADNI datasets (ADNI-1 and ADNI-2).

### 3.1 SNP-based analysis

From ADNI-1 dataset analysis, chromosome 20 and in particular the 20orf196 gene appeared relevant in the AD context. Although the genes and SNPs located in the locus 20p12.3, are not included in known linkage regions associated to AD, according to AlzGene database (Bertram *et al.*, 2007), the identification of SNPs nearby the same gene in both binary classification tasks (SNP **rs6053572** mapped on the intergenic region GPCPD1-20orf196 in cases@controls; and SNPs **rs236137** and **rs6041271** on the intergenic region 20orf196-CHGB in APOEε4 task) strengthened its implication in AD. In addition, within the ADNI-2 dataset, the gene LOC101928063 and its SNP **rs367209** on chromosome 19 was strongly associated to AD cases@controls and APOEε4 task (Table S2). Chromosome 19 was also in common with ADNI-1 SNP signature, considering the APOEε4 task, although the related SNPs identified harbored on different genes and intergenic regions.

Considering that the demographic and clinical characteristics of the subjects enrolled in the two studies were similar, a possible explanation of the low reproducibility between the two SNP signatures could be due to the different Illumina GWAS platforms used. ADNI-1 and ADNI-2 datasets reported 620901 and 730523 SNPs respectively, of which only 300000 were in common. Therefore, the use of different platforms might be a critical point in the context of data validation. On the other hand, the functional characterization of the ADNI-1 and ADNI-2 SNP signatures found enriched the same KEGG pathway “**Neuroactive ligand-receptor interaction**” in which some genes encoded for glutamate receptors were found in both datasets. For example the GRM7 gene, that encodes for the glutamate metabotropic receptor 7 and plays an important role as neurotransmitter in the cerebral cortex, hippocampus, and cerebellum (Makoff *et al.*, 1996). Epidemiologic studies have identified associations between variation in *GRM7* and depression, anxiety, schizophrenia, bipolar disorder, and epilepsy (Haenisch *et al.*, 2015; Chen *et al.*, 2018). Recently it has also been demonstrated that 3xTg-AD mice showed lower GRM7 protein expression in hippocampus, associated with an increased anxiety behavior, compared with the wild-type mice (Zhang *et al.*, 2016). The significance of such results was confirmed by a genome-wide gene and pathway-based analyses on depressive symptom burden in the three independent cohort (Nho *et al.*, 2015). Genes encoding for NMDA glutamate ionotropic receptor subunits, the major ion channel that participates in neuronal development and synaptic plasticity (Kehoe, Bernardinelli and Muller, 2013) were also identified in the two datasets. In particular, the GRIN3A, found in ADNI-1, is located within 9q21.31-q32, that is linkage region associated to AD (Bertram *et al.*, 2007). Deregulations of GRIN3A activity has been associated with abnormalities in dendritic spine density, turnover, formation and elimination observed in different neurological conditions, including AD (Kehoe, Bernardinelli and Muller, 2013). While, GRIN2A, interacting with GRIN3A, are globally involved in the modulation of episodic memory consolidation (Papenberg *et al.*, 2014) and of the response to antipsychotic treatment (Stevenson *et al.*, 2016).

Others genes of ADNI-1 and ADNI-2 signature, such as P2RY13, P2RY14, GABRG3 associated to the “Neuroactive ligand-receptor interaction” pathway, might also have a relevance in AD pathology. For example two members of the G-protein coupled receptors family named P2RY13 and P2RY14: the first gene is known to be involved in the axonal elongation (del Puerto *et al.*, 2012), while the second gene has been reported to play a role in neuroimmune function (Sesma *et al.*, 2012; Barrett *et al.*, 2013). Both these genes are located within 3q12.3-q25.31, a linkage region known to be associated to AD (Bertram *et al.*, 2007). Furthermore, gamma3-aminobutyric acid (GABA) receptor, encoded by GABRG3 gene, could have a relevance in protecting neurons against neurofibrillary pathology in AD (Iwakiri *et al.*, 2009).

### 3.2 Gene-based analysis

Surprising, a gene signature was successfully found only for ADNI-1. Again, this diff erence in ADNI-1 and ADNI-2 results could be due to the low overlap of SNP measured in the two different platforms.

The gene-based analysis of ADNI-1 dataset confirmed the association of TOMM40 gene with AD pathology, according with different other studies (Linnertz *et al.*, 2014; Goh *et al.*, 2015; Chiba-Falek, Gottschalk and Lutz, 2018). In particular, TOMM40, with its three SNPs was found in common between cases@controls and APOEε4 tasks. TOMM40 is located in 19q13.32 locus, a known linkage region for AD (Bertram *et al.*, 2007). Its encoded protein plays a key role in the mitochondria functionality being essential for import of protein precursors into mitochondria. Furthermore, one of the three SNPs mapped to this gene, rs2075650, is also known to be a contributing factor for AD (Huang *et al.*, 2016) (Potkin *et al.*, 2009). In addition, in the cases@controls task, SKAT analysis identified also TEF gene, a member of the PAR subfamily of the basic region/leucine zipper (bZIP) family. The rs738499 SNP mapped to TEF and identified in ADNI-1, has been found to be associated to sleep disturbances in Parkinson’s (Hua *et al.*, 2012) and in Alzheimer’s diseases (Gast *et al.*, 2012).

Interestingly, comparing the SNP and the gene signatures, the TOMM40 gene with its two SNPs, rs2075650 and rs8106922, and the intergenic region LOC100129500-APOC1 with rs439401 SNP (Fig. S2) were found in common with the two signatures.

#### 3.3 Pathway-based analysis

Many statistically significant pathways were found in common between ADNI-1 and ADNI-2. Specifically for APOEε4 task, the common groups in the two datasets were **5a, 1c and 9a**, related to the metabolism of proteins (*e.g.*, amyloid fiber formation, unfolded protein response, and mitochondrial protein import), to the cellular response to stress (*e.g.*, detoxification of reactive oxygen species) and to the signaling by NOTCH and GPCR pathways respectively (Table 2). It is noteworthy that the “mitochondrial protein import” pathway, related to group 5a, in common with both datasets, involved TOMM40 gene, resulting also from SNP and gene-base analysis of ADNI-1 dataset. A pathway in common to both datasets, belonging to group 9a was “GPCR ligand binding” in which GRM7, and the GRIN2A and GRIN3A (the two NMDA glutamate receptor subunits) were involved.

In conclusion, this study pointed out a promising approach to get more insight in studying heritable susceptibility of a complex disease like AD. What has been raised by this approach was the difficulty to reproduce and validate the specific signatures on different datasets, since the validation procedure truly succeeded just for the APOEε4 signature identified in ADNI-1 (Fig. S2). Nevertheless, considering the **telescope** approach with multivariate analysis, some promising candidate, such as TOMM40 and GRM7, was confirmed in the two datasets. In future, this list might be increased analyzing additional GWAS datasets, thus contributing to obtain a robust global signature of AD susceptibility.

## 4. Star Methods

### 4.1 Datasets

Data used in the preparation of this article were obtained from the Alzheimer’s Disease Neuroimaging Initiative (ADNI) database (adni.loni.usc.edu). The primary goal of ADNI has been to test whether serial magnetic resonance imaging (MRI), positron emission tomography (PET), other biological markers, and clinical and neuropsychological assessment can be combined to measure the progression of mild cognitive impairment (MCI) and early AD. Further information about ADNI can be found here (Weiner *et al.*, 2010) (http://www.adni-info.org). In this study Genome-wide association studies (GWAS) data and APOE genotype obtained in the ADNI-1 and ADNI-2 datasets (Saykin *et al.*, 2010) were used (Table S1), considering the AD and healthy controls (CN) group. The genotyping platforms used by ADNI-1 and ADNI-2 were: Illumina Human 610-Quad BeadChip that measures 620.901 SNPs and CNV markers for ADNI-1 and Illumina Human OminExpress-24v that measured 730.525 SNPs and CNV markers for ADNI-2. Differently from ADNI-2, in ADNI-1 APOE genotyping is provided outside the GWAS platform. In both datasets, we performed two supervised binary classification analyses: AD vs. cognitively healthy subjects (cases@controls task) and subjects at risk vs. subjects not at risk of developing AD, according with APOE status (1 or 2 alleles vs. 0 allele of APOEε4) (APOEε4 task) (Table S1).

### 4.2 Workflow

A 3-fold kernel approach was devised to analyze the datasets. Considering the two classification tasks addressed and the 3-fold analyses, each ADNI dataset was analyzed six times (Fig. 1).

In order to increase the signal over noise ratio, reducing the number of SNPs to analyze, we adopted the following strategy: (1) for the SNP and pathway-based analysis we employed two sparse methods, designed to identify the SNPs or pathways which are most discriminative for the classification tasks while restricting the selection of SNPs and pathways, and considered a different representation of the SNP data (see Supplementary Materials); (2) for the SNP-based analysis we analyzed each chromosome separately while for the gene and pathway-based analyses we grouped the SNPs considering genes/intergenic regions or pathways relevant for AD.

### 4.3 SNP-based analysis

For the SNP-based analysis, l1l2 feature selection (l_1_l_2FS_), a method that belong to sparse techniques was chosen (Hastie *et al.*, 2015). This method allows the identification of the most discriminative variables for the problem at hand (classification tasks) while making feature selection (see Supplementary Materials). l l was used within PALLADIO (https://github.com/slipguru/palladio), a machine learning Python library that can be customized to consider various combinations of feature selections and classification methods (Fig. S1A). In order to ensure the reliability of the results, we used PALLADIO to perform two sets of experiments, which we referred to as *regular* batch and *permutation* batch (Fig. S1B). The level of distance of the two distributions measured the reliability of the obtained results: the higher the distance, more reliable are the obtained results (see Supplementary Materials).

### 4.4 Gene-based analysis

For the gene-based analysis, three different association tests available in the SKAT package were used: Burden, SKAT and SKATO (see Supplementary Methods). SKAT (Wu *et al.*, 2011; Ionita-Laza *et al.*, 2013) is a supervised regression method that test the association between genetic variants in a region and a dichotomous or a continuous trait while adjusting for covariates. The dichotomous traits considered were cases@controls and APOEε4 task. Covariates such as age at onset, race, sex were excluded from the analysis. Furthermore we chose to consider genes or intergenic regions, leveraging on the mapping files SNPs-to-genes provided by the GWAS platform manufacturer (i.e., “Human610_Gene_Annotation_hg19.txt” for ADNI-1 and “HumanOmniExpress-24v1-1_Annotated.txt” for ADNI-2).

The threshold of genome-wide significance we established, was in accordance with other studies (Kraft, Zeggini and Ioannidis, 2009; Mukherjee *et al.*, 2014; Fadista *et al.*, 2016; Kanai, Tanaka and Okada, 2016) (see Supplementary Methods).

### 4.5 Pathway-based analysis

We selected 9 groups of pathways more relevant for neurodegenerative processes (Table S2) inside REACTOME database (Croft *et al.*, 2013). Each group contained two or more pathways and each group represented a SNP matrix that, together with a label that characterizes each subject, was given as input to “Group Lasso with overlap” (Hastie *et al.*, 2015). This latter is a machine learning method, able to consider the presence of overlapping groups of SNPs mapped to genes, involved in more than one pathway inside a group. The goal of “Group Lasso with overlap” is to induce a “sparse” selection at the group level, using all the pathways specified in the group. In this way, starting from a possibly long list of pathways inside a group, the algorithm selected a few (but informative) pathways that could be relevant for the problem at hand.

## Acknowledgements

Data collection and sharing for this project was funded by the Alzheimer’s Disease Neu ro imaging Initiative (ADNI) (National Institutes of Health Grant U01 AG024904) and DOD ADNI (Department of Defense award number W81XWH-12-2-0012). ADNI is funded by the National Institute on Aging, the National Institute of Biomedical Imaging and Bioengineering, and through generous contributions from the following: AbbVie, Alzheimer’s Association; Alzheimer’s Drug Discovery Foundation; Araclon Biotech; BioClinica, Inc.; Biogen; Bristol-Myers Squibb Company; CereSpir, Inc.; Cogstate; Eisai Inc.; Elan Pharmaceuticals, Inc.; Eli Lilly and Company; EuroImmun; F. Hoffmann-La Roche Ltd and its affiliated company Genentech, Inc.; Fujirebio; GE Healthcare; IXICO Ltd.; Janssen Alzheimer Immunotherapy Research & Development, LLC.; Johnson & Johnson Pharmaceutical Research & Development LLC.; Lumosity; Lundbeck; Merck & Co., Inc.; Meso Scale Diagnostics, LLC.; NeuroRx Research; Neurotrack Technologies; Novartis Pharmaceuticals Corporation; Pfizer Inc.; Piramal Imaging; Servier; Takeda Pharmaceutical Company; and Transition Therapeutics. The Canadian Institutes of Health Research is providing funds to support ADNI clinical sites in Canada. Private sector contributions are facilitated by the Foundation for the National Institutes of Health (www.fnih.org). The grantee organization is the Northern California Institute for Research and Education, and the study is coordinated by the Alzheimer’s Therapeutic Research Institute at the University of Southern California. ADNI data are disseminated by the Laboratory for Neuro Imaging at the University of Southern California.

## Author contributions

M.S. contributed to conception and design of the study; analysis and interpretation of the data; preparation of the final manuscript. F. T. and V. T. contributed to the implementation of the method Group Lasso with overlap used in the Pathway-based analysis. A. B. contributed to the revision of the final manuscript. D. U. contributed to the interpretation of the data, to the preparation and revision of the final manuscript.

## Supplemental Information

Document S1. It contains further explanations of the methods utilized and the results of the validation of the SNP signatures. It also contains Figures S1, S2 and Tables S1-S4.

## Figure/Tables legends

**Table 1. Functional characterization of ADNI-1 and ADNI-2 SNP signatures**. This analysis was focused on the gene lists derived from the ADNI-1 SNP signature (APOEε4 task) and from the ADNI-2 SNP signature (cases@control task). The number of genes, their gene symbols and the Adjusted P value (Adj-P value) is reported for each significant pathway or disease found enriched. The significance level chosen for the enrichment analysis was .05.

**Table 2**. **Gene-based signatures identified in ADNI-1**. Lists of genes identified by the SKAT software in the cases@controls and APOEε4 tasks. The genes with P value < 1.37×10^−6^ are considered significant.

**Table 3**. **Pathway-based signatures identified in ADNI-1 and ADNI-2**. Lists of the groups of pathways found statistically significant in APOEε4 task for ADNI-1 and in both tasks (cases@controls and APOEε4) for ADNI-2. The groups 1c, 5a, 9a were in common with ADNI-1 and 2. The test score shows the classification performance of “Group Lasso with overlap”. See Table S2 for the complete list of all the pathways analyzed inside each group.

